# CDK4 as a phytochemical based anticancer drug target

**DOI:** 10.1101/859595

**Authors:** Rony Kumer Chando, Nazmul Hussain, Mashedul Islam Rana, Shomoita Sayed, Saruar Alam, Tawkir Ahmed Fakir, Sudip Sharma, Arifur Rahman Tanu, Faraid Mobin, Ehsanul Hoque Apu, Kamrul Hasan, Abu Sayed, Mohammad Arif Ashraf

**Affiliations:** Department of Biochemistry and Molecular Biology, Tejgaon College, 16 Indira Rd, Dhaka - 1215, Bangladesh; Department of Biochemistry and Molecular Biology, Hajee Mohammad Danesh Science and Technology University, Dinajpur-5200, Bangladesh; Department of Biochemistry and Molecular Biology, University of Dhaka, Dhaka, Bangladesh; Technical University of Munich, 80333 München, Germany; Institue for Genomics and Evolutionary Medicine (iGEM), Temple University, Philadelphia, PA, USA; Department of Biology, Temple University, Philadelphia, PA 19122, USA; Ministry of Labour and Employment, People’s Republic of Bangladesh, Dhaka, Bangladesh; Institute for Quantitative Health Science and Engineering (IQ), Department of Biomedical Engineering, Michigan State University, East Lansing, MI 48824, USA; Biology department, University of Massachusetts Amherst, Amherst, MA 01002, USA

## Abstract

Success story of plant-based medicine had been overlooked during the advent of modern pharmaceutical industry. Despite the negligence of the multimillion-dollar drug industry, people entirely rely on medicinal plants in some part of the world. In this study, we have emphasized on going back to those traditional medicinal practices to figure out their underlying mechanism to move forward on phytochemical based drug development. We screened Medicinal Plant Database Bangladesh 1.0 (MPDB1.0) and on-going extension, MPDB2.0, of that database to find traditionally used medicinal plants and their active compounds. Here, Mangiferin, extracted from *Mangifera indica*, have been demonstrated to interact with cell cycle regulator Cyclin-dependent Kinase 4 (CDK4). *CDK4* is differentially expressed during Glioblastoma multiforme (GBM), Brain Lower Grade Glioma (LGG), and Sarcoma (SARC). Expression of *CDK4* is interlinked to the patients’ survival rate and its consistent expression throughout different stages have provided the advantage to use it as diagnostic tool and drug target. Altogether, this study demonstrated that simple mango tree extracted active compounds, mangiferin, can work as potential anticancer drug and leveraging the recent advancement of sequencing and gene expression data can accelerate the phytochemical based drug discovery process.

## Introduction

Cancer is expected to supersede all other non-communicable diseases to become the major cause of death and the most indomitable barrier to increasing life expectancy in every country of the world in the 21st century (Bray, Ferlay, and Soerjomataram 2018). World Health Organization (WHO) in 2015 estimated cancer to be the first leading cause of death before age 70 years in 91 of 172 countries, and also appraised it as the third or fourth major cause in an additional 22 countries. According to the GLOBOCAN 2018 estimation, 18.1 million new cancer cases and 9.6 million cancer deaths occurred in 2018 among which lung cancer is the most commonly diagnosed cancer (11.6% of the total cases) closely followed by female breast cancer (11.6%), prostate cancer (7.1%), and colorectal cancer (6.1%) for incidence (Bray, Ferlay, and Soerjomataram 2018). Approximate number of cancer patients in Bangladesh vacillates between 1.3 to 1.5 million, with about an addition of 0.2 million new patients each year (Uddin et al. 2016; Noronha et al. 2012). Despite of being a compilation of approximate 5 decades of systemic drug delivery and establishment, a repertoire of chemotherapeutic drugs which is the standard cancer treatment is not bereft of their own intrinsic problems such as toxicity and lesser efficacy (Desai et al. 2008). In the last decade, identification of medicinal plants with significant cytotoxic potential useful for the development of cancer therapeutics has become a center of attention with still lots of unexplored areas for elucidation via research (Al-kalaldeh, Abu-dahab, and Afifi 2010). More than 1000 plants species have been identified with significant anticancer potential (Mukherjee et al. 2001). The isolation of the vinca alkaloids, vinblastine (Balunas and Kinghorn 2005) from the Madagascar periwinkle, Catharanthus roseus G. Don. (Apocynaceae) bolstered medicinal plants utilization as a promising source of anti-cancer medication. This in combination with vincristine and other cancer chemotherapeutic drugs are used for the treatment of a spectrum of cancers such as leukemias, lymphomas, advanced testicular cancer, breast and lung cancers, and Kaposi’s sarcoma (Cragg and Newman 2005). The discovery of paclitaxel (Taxol) (Butler 2004) from the bark of the Pacific Yew, Taxus brevifolia Nutt. (Taxaceae), is another evidence of the success in natural product drug discovery. Utilization of various parts of Taxus brevifolia from which paclitaxel was discovered and other Taxus species (e.g., Taxus Canadensis Marshall, Taxus baccata L.) by several Native American Tribes sheds light on indigenous knowledge of medicinal plants (Cragg and Newman 2005). Another potent plant-acquired active compound, Homoharringtonine (Norman et al. 1985) was isolated from the Chinese tree *Cephalotaxus harringtonia* var. drupacea (Sieb and Zucc.) (Cephalotaxaceae) and has been used successfully in China in a racemic mixture with harringtonine for the treatment of acute myelogenous leukemia (Cragg and Newman 2005). Elliptinium, a derivative of ellipticine, isolated from a Fijian medicinal plant Bleekeria vitensis A.C. Sm., is marketed in France for the treatment of breast cancer (Cragg and Newman, 2005).

With a distinguished heritage of herbal medicines for primary health care among the South Asian countries, Bangladesh is estimably home to more than thousands of species of medicinal plants. These native plants are a considerable source of Unani, Ayurvedic and homeopathic medicines in Bangladesh (Gani et al., 2003) with their background entrenched in folklores and century-old knowledge of traditional medicine practitioners (Mohammed et al. 2010).

Unfortunately, major portion of these plants have not yet been studied extensively in terms of their chemical, pharmacological and toxicological properties to explore their bioactive compounds which may prove to be an astonishing source of new anticancer drug discovery (Khatun et al. 2014). A comprehensive database including all endemic medicinal plants works as a foundational basis for future drug discovery. There have been such extensive, curated databases such IPPAT (Indian Medicinal Plants, Phytochemistry And Therapeutics) consisting 1742 Indian Medicinal Plants, 9596 Phytochemicals and 1124 Therapeutic uses spanning 27074 plant-phytochemical associations (Mohanraj, Karthikeyan, Chand, et al. 2018). CMKb (Customary Medicinal Knowledge) is another such endeavor for storing, preserving and circulating aboriginal Australian medicinal plant knowledge (Gaikwad et al. 2008). In this article, we are mainly shedding light on changes in several anti-neoplasmic bioactive compounds found in Bangladeshi indigenous plants when they are bound to proteins of cancer cascade pathways with cheminformatic approaches.

We have identified Mangiferin, a xanthonoid extracted from the bark and leaves of mango tree (Mangifera indica), as a potential anticancer drug which targets cell cycle Cyclin-dependent kinase 4 (CDK4). This study took the leverage of curated medicinal plants’ information of Bangladesh to screen potential anticancer drugs from phytochemicals and used the large scale RNA sequencing data to analyze expression of CDK4 in cancer sub-types, different stages of cancer, survival events correlated with the expression, and co-expression as well. This study is a big step forward to broaden our understanding about the primitive plant-based medicine and bridging this ancient knowledge with cutting-edge large-scale cancer dataset.

## Materials and Methods

### Database creation

MPDB 2.0 is the continuation of the MPDB 1.0 (http://www.medicinalplantbd.net/) which contains the information such as scientific name, family name, local name, utilized part, location, ailment, active compounds and PubMed ID of related research article about medicinal plants from Bangladeshi. To acquire this information regarding medicinal plants from Bangladesh, ∼75 research, survey, and review articles (published in both national and international journals till September 2019) were considered. As most of these articles lack the knowledge about active ingredients, we have used the scientific name of these plants to search one more round through PubMed to find out reported active compounds extracted from these plants.

### Identification of interacting proteins

The interaction between plant-based active compounds and human proteins was investigated by using STITCH 4.0 (http://stitch.embl.de/); a web server focusing on the interactions between proteins and small molecules. STITCH (Search Tool for Interactions of Chemicals) amalgamates information about such interactions from metabolic pathways, crystal structures, binding experiments and drug–target relationships (Kuhn et al. 2008).

### Molecular docking

3-D structure of the proteins are derived from RCSB PDB (http://rcsb.org). RCSB PDB is the single global archive for experimentally determined, atomic-level three dimensional structures of biological macromolecules in PDB format (Rose *et al.*, 2017). Chemical structures of active compounds are attained from PubChem (https://pubchem.ncbi.nlm.nih.gov), a public repository for information on chemical substances and their biological activities in (S. Kim et al. 2016; Wang et al. 2010; 2012; 2009). The phytochemical compounds are docked against the proteins using PyRx (https://pyrx.sourceforge.io/), an open-source software with virtual molecular screening ability to dock small-molecule libraries to a macromolecule with an aim to discover lead compounds with desired biological function (Dallakyan and Olson 2015). For the docking of targeted phytochemicals into discovered protein binding pockets (Sousa, Fernandes, and Joa 2006) and to approximate the binding affinities of docked ligands, a molecular docking program AutoDock Vina (Oleg and J. 2011) in PyRx Virtual screening tool is mainly employed. The protein PDB file was changed into the PDBQT format file containing the protein atom coordinates, partial charges and deliverance parameters and the ligands file (SDF) are distorted into PDBQT format (Saddala et al. 2016).

### Large-scale gene expression analysis

Over the last few years, cancer related large scale RNA sequencing data has been available through TCGA and GTEx (Consortium 2015; Lonsdale et al. 2013; Weinstein et al. 2013). This data has become more accessible through GEPIA (Gene Expression Profiling Interactive Analysis) (Tang et al. 2017) and their recent updated GEPIA2 (Tang et al. 2019). For gene expression analysis in cancer sub-types, differential methods LIMMA was used with log2FC cutoff 1 and q-value cutoff 0.01. In stage-wise expression analysis, major stages were only considered. During the patients’ survival analysis, data was normalized using overall survival with 95% confidence interval. Correlation analysis is based on Pearson correlation test.

### Statistical analysis and graphs

Statistical analysis and graphs are either generated by web server as mentioned in the manuscript or other cases, used statistical programming language R (3.6.0). R codes are available from the authors upon request. Final figures were prepared on Adobe Illustrator (version 24.0.1) without compromising the details of the analysis.

## Results

### Anticancer compounds are prevalent among traditional plant extracts

We have started to gather the information of traditionally used medicinal plants of Bangladesh based on Medicinal Plant Database Bangladesh (MPDB1.0) (Ashraf et al. 2014), which contains 353 plants and known active compounds for 78 plants. This search was extended, and 2,349 new plants information included along with known active compounds is considered for this study. The extended dataset is considered and mentioned as MPDB 2.0 (unpublished) in this manuscript. Among 2,702 traditionally used plants, we have searched published active compounds reported for their role in regulating diabetes, respiratory disorder, jaundice, cancer, diarrhea, skin disease and so on. Interestingly, we have observed that highest number of active compounds, 42, are reported for the anticancer activity (Figure 1a). Unfortunately, majority of these compounds are neither used for clinical trial nor the anticancer mechanism is known.

**Figure 1:**
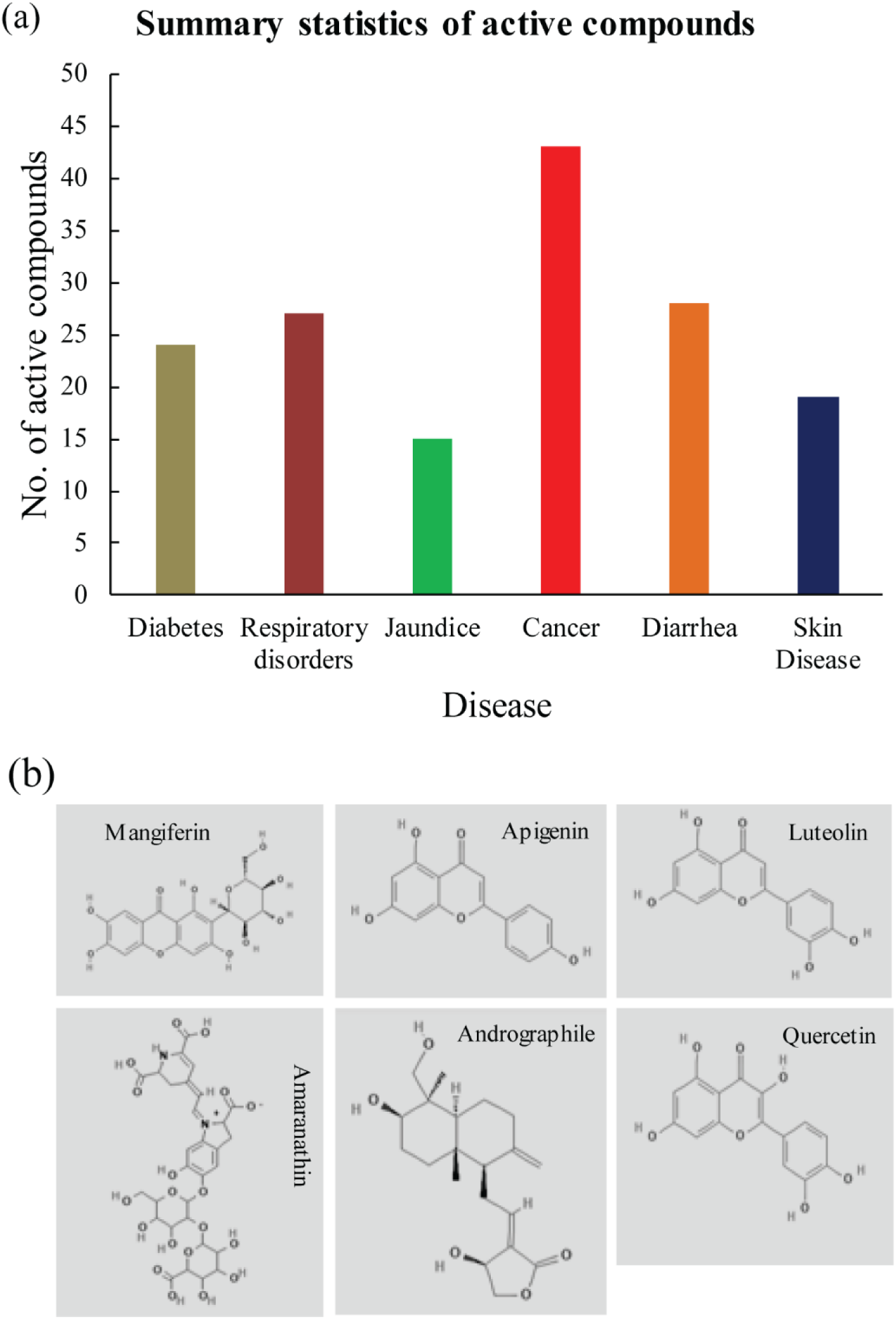
(a) Summary statistics of identified active compounds based on their known roles from literature and traditional medicinal practice. (b) Selected chemicals from the cancer target identification. From top – left: Mangiferin, Apigenin, Luteolin and from bottom – left: Amaranthin, Andrographile, Quercetin.

### Cell cycle regulatory genes are targeted by identified active compounds

Using the identified active compounds reported for cancer (Figure 1a), we tried to find their interacting human proteins. Out of 42 active compounds, we have found interacting protein hits for 12 compounds (tannic acid, quercetin, betacyanin, amaranthin, mangiferin, voacangine, quinic acid, andrographolide, luteolin, apigenin, rutin, gallic acid). Among these compounds, tannic acid (Bridgeman, Nguyen, and Kishore 2018), quercetin (J. Jeong et al. 2009), mangiferin (Núñez Selles, Daglia, and Rastrelli 2016), voacangine (Y. Kim, Jung, and Kwon 2012), quinic acid (Singh, Chauhan, and Tripathi 2018), andrographolide (Peng et al. 2018), luteolin (Cook 2018), apigenin (Yan et al. 2017), rutin (Khan et al. 2019), and gallic acid (Liu et al. 2012) are already reported for their anticancer activities. This suggests that the screening process from the traditionally used medicinal plants contain both potential anticancer properties.

Against these 12 compounds, we could identify 83 interacting protein targets. These targeted proteins are mostly enriched by cell cycle regulator proteins. We have selected 6 active compounds (andrographolide, luteolin, apigenin, quercetin, amaranthin, and mangiferin) (Figure 1b) and 7 interacting proteins (CCR3, CDK2, MAPK8, TP53, HIBCH, NOS3, and CDK4) to test their binding affinity through molecular docking study. Binding affinity ranges from −7.8 kcal/mol to - 9.6 kcal/mol (Figure 2, Table 1). In general, the most negative numerical value for the binding affinity indicates the best predicted binding between a ligand and a macromolecule (Dallakyan and Olson 2015). This result indicates that these plant-based active compounds have potential binding affinity with identified protein targets.

**Table 1:** Details of molecular docking

| Ligand | Macromolecule | Binding<br>affinity<br>(Kcal/mol) | X | Y | Z |
| --- | --- | --- | --- | --- | --- |
| Mangiferin | CDK4 | -7.8 | 69.0904 | 50.5133 | 62.8484 |
| Quercetin | HBICH | -8.6 | 48.1841 | 65.0171 | 64.423 |
| Andrographodile | CCR3 | -9.3 | 69.5242 | 66.1132 | 70.1861 |
| Luteolin | CDK2 | -8.8 | 55.8984 | 39.4384 | 64.6451 |
| Luteolin | MAPK8 | -9.2 | 61.7916 | 67.581 | 68.3343 |
| Amaranthin | NOS3 | -9.6 | 60.2027 | 63.829 | 75.0564 |
| Apigenin | TP53 | -7.9 | 51.7768 | 48.08834 | 62.6262 |

**Figure 2:**
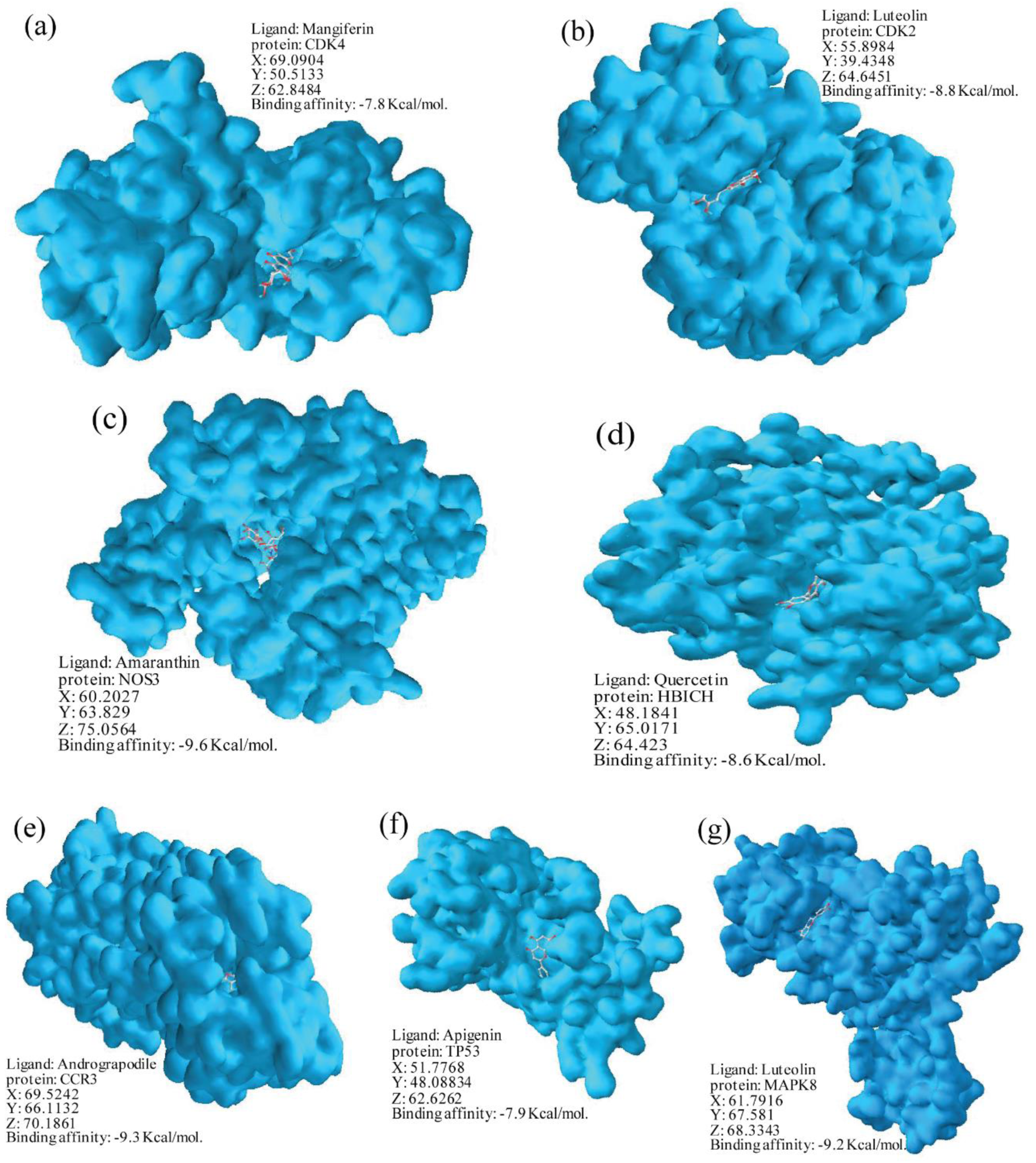
Binding affinity assay between active compounds and proteins. Docking combination of (a) Mangiferin – CDK4; (b) Luteolin – CDK2; (c) Amaranthin – NOS3; (d) Quercetin – HBICH; (e) Andrograpodile – CCR3; (f) Apigenin – TP53; (g) Luteolin – MAPK8 are presented with binding affinity (Kcal/mol) and x, y, z coordinates used for the docking.

### CDK4 expression is differentially regulated by multiple cancer

The combination of mangiferin and Cyclin-dependent kinase 4 (CDK4) seems interesting because CDK4 inhibition is potential target for cancer treatment (Goel et al. 2018). As a fist step of targeting the CDK4, we tried to find out in which type of cancer *CDK4* is differentially expressed. We have analyzed the gene expression data of *CDK4* in 33 cancer sub-types: ACC (Adrenocortical carcinoma), BLCA (Bladder Urothelia Carcinoma), BRCA (Breast invasive carcinoma), CESC (Cervical squamous cell carcinoma and endocervical adenocarcinoma), CHOL (Cholangio carcinoma), COAD (Colon adenocarcinoma), DLBC (Lymphoid Neoplasm Diffuse Large B-cell Lymphoma), ESCA (Esophageal carcinoma), GBM (Glioblastoma multiforme), HNSC (Head and Neck squamous cell carcinoma), KICK (Kidney Chromophobe), KIRC (Kidney Renal clear cell carcinoma), KIRP (Kidney renal papillary cell carcinoma), LAML (Acute Myeloid Leukemia), LGG (Brain Lower Grade Glioma), LIHC (Liver hepatocellular carcinoma), LUAD (Lung adenocarcinoma), LUSC (Lung squamous cell carcinoma), MESO (Mesothelioma), OV (Ovarian serous cystadenocarcinoma), PAAD (Pancreatic adenocarcinoma), PCPG (Pheochromocytoma and Paraganglioma), PRAD (Prostate adenocarcinoma), READ (Rectum adenocarcinoma), SARC (Sarcoma), SKMC (Skin Cutaneous Melanoma), STAD (Stomach adenocarcinoma), TGCT (Testicular Germ Cell Tumors), THCA (Thyroid carcinoma), THYM (Thymoma), UCEC (Uterine Corpus Endometrial Carcinoma), UCS (Uterine Carcinosarcoma), UVM (Uveal Melanoma) (Figure 3). *CDK4* is highly expressed in GBM, LGG, and SARC (Figure 3). This observation is further tested by comparing the *CDK4* expression between tumor and normal cells in GBM, LGG, and SARC. *CDK4* expression differs significantly in GBM and LGG, but the difference in SARC is not significant due to lack of enough normal cell data (Figure 4). Additionally, expression of *CDK4* is analyzed at different stage of GBM, LGG, and SARC (Figure 5). Its expression is consistent and higher in every stage compared to *TP53* indicates the potential use of *CDK4* expression for diagnostic and drug target candidate (Figure 5a, 5b). *CCR3* expression is used as negative control (Figure 5c). This gene expression analysis clearly indicates that *CDK4* expression is a prime indicator of uncontrolled cell division during cancerous growth and their expression compared to *TP53* strengthen the evidence.

**Figure 3:**
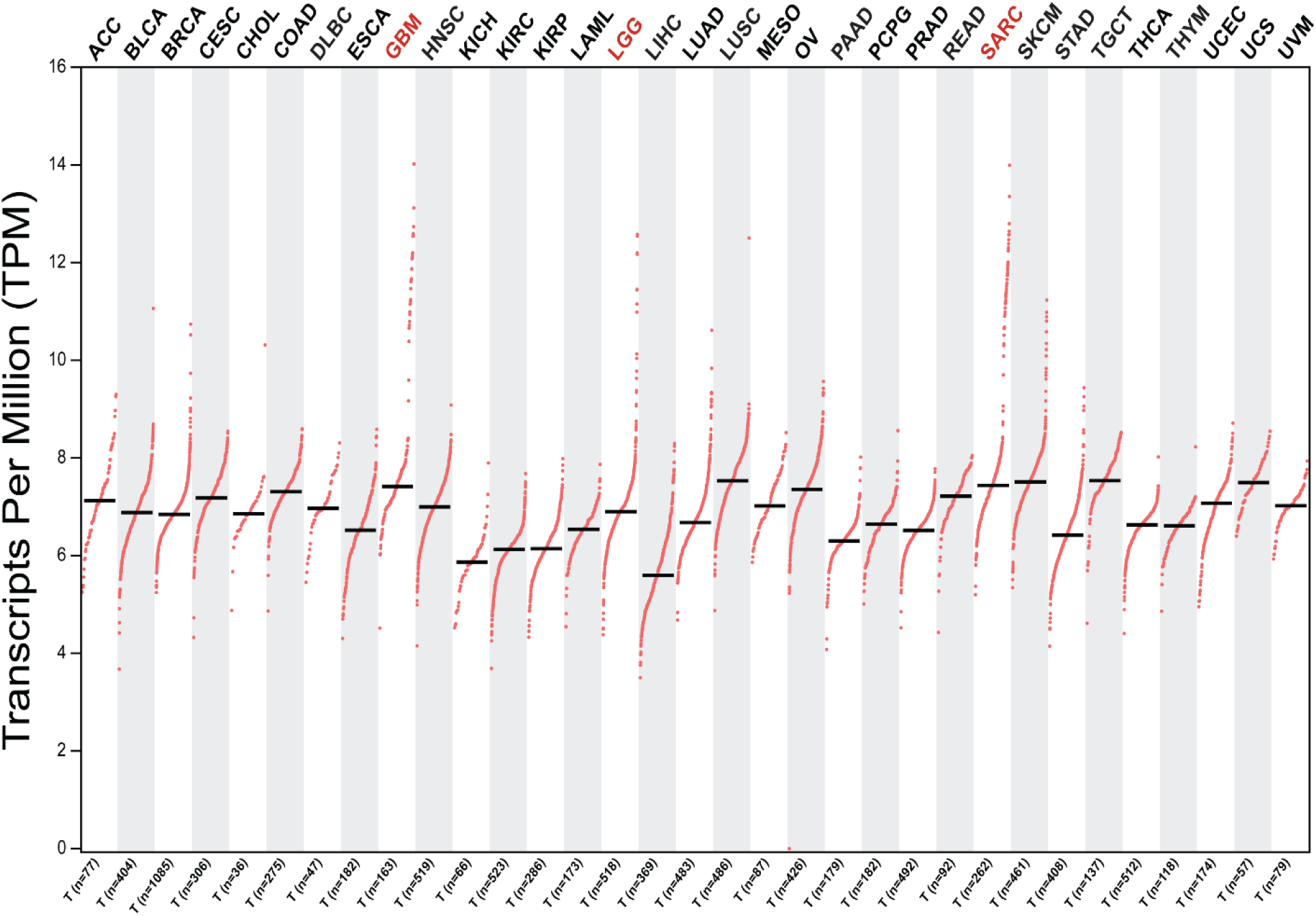
*CDK4* expression analysis. *CDK4* expression is demonstrated in ACC (Adrenocortical carcinoma), BLCA (Bladder Urothelia Carcinoma), BRCA (Breast invasive carcinoma), CESC (Cervical squamous cell carcinoma and endocervical adenocarcinoma), CHOL (Cholangio carcinoma), COAD (Colon adenocarcinoma), DLBC (Lymphoid Neoplasm Diffuse Large B-cell Lymphoma), ESCA (Esophageal carcinoma), GBM (Glioblastoma multiforme), HNSC (Head and Neck squamous cell carcinoma), KICK (Kidney Chromophobe), KIRC (Kidney Renal clear cell carcinoma), KIRP (Kidney renal papillary cell carcinoma), LAML (Acute Myeloid Leukemia), LGG (Brain Lower Grade Glioma), LIHC (Liver hepatocellular carcinoma), LUAD (Lung adenocarcinoma), LUSC (Lung squamous cell carcinoma), MESO (Mesothelioma), OV (Ovarian serous cystadenocarcinoma), PAAD (Pancreatic adenocarcinoma), PCPG (Pheochromocytoma and Paraganglioma), PRAD (Prostate adenocarcinoma), READ (Rectum adenocarcinoma), SARC (Sarcoma), SKMC (Skin Cutaneous Melanoma), STAD (Stomach adenocarcinoma), TGCT (Testicular Germ Cell Tumors), THCA (Thyroid carcinoma), THYM (Thymoma), UCEC (Uterine Corpus Endometrial Carcinoma), UCS (Uterine Carcinosarcoma), UVM (Uveal Melanoma).

**Figure 4:**
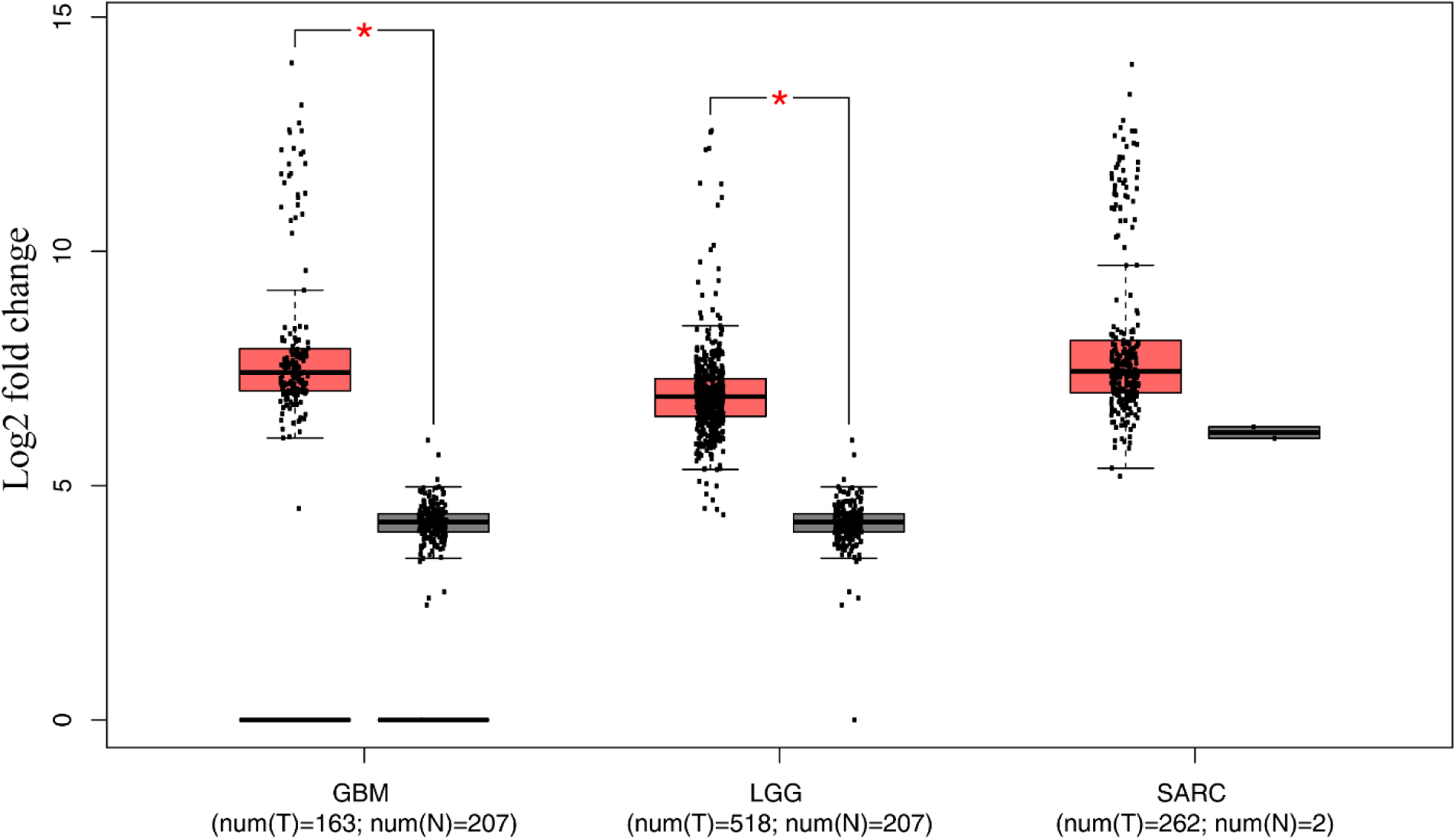
*CDK4* expression in GBM, LGG, and SARC. *CDK4* expression is demonstrated for GGM, LGG, SARC between cancerous and normal cells. Expression was considered in Log2 fold change and data points used here are mentioned for type of cancer.

**Figure 5:**
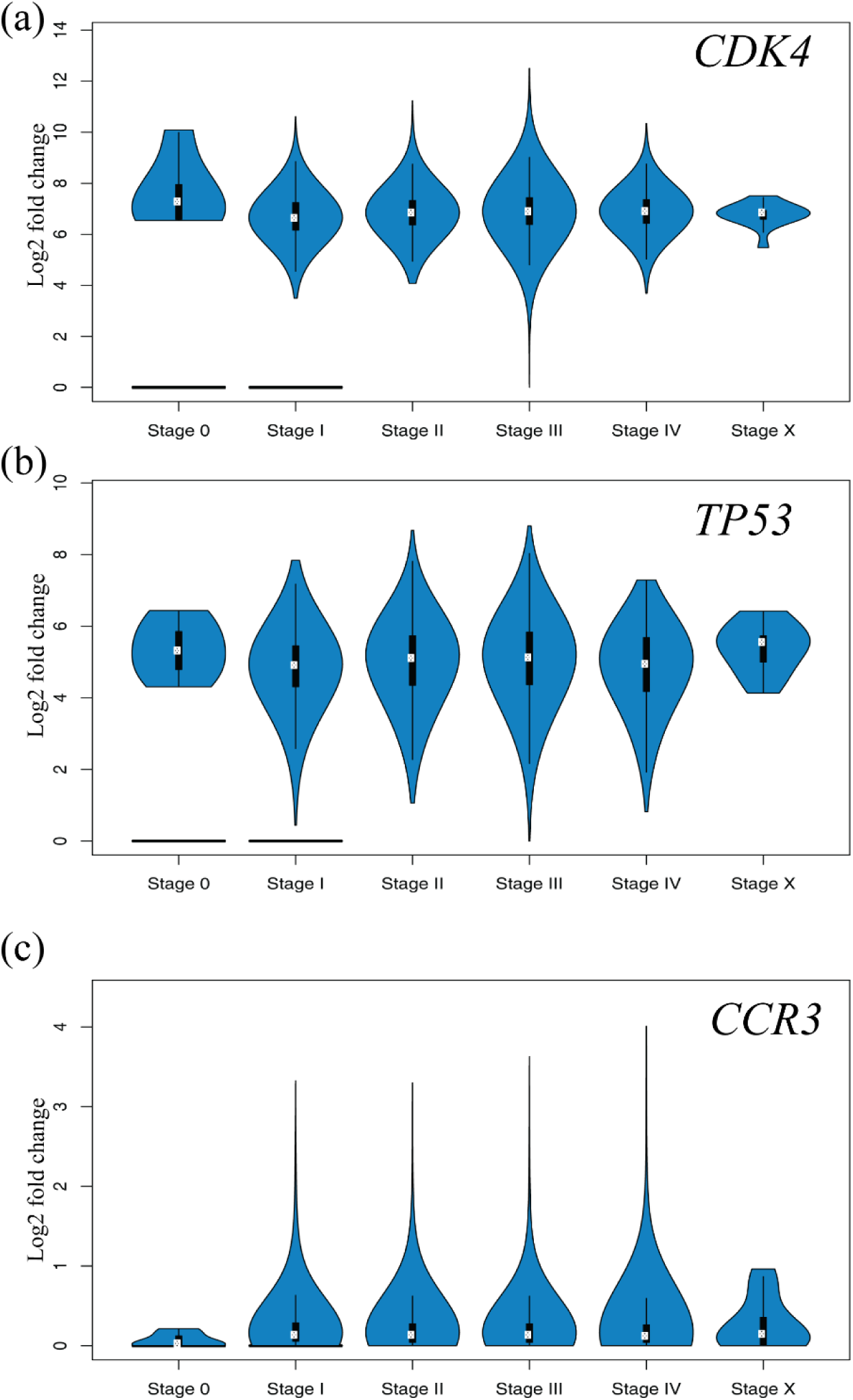
Gene expression analysis at different stages of cancer. Expression of *CDK4* (a), *TP53* (b), and *CCR3* (c) at different stages of multiple cancers are provided in log2 fold change scale.

### CDK4 as potential mangiferin based drug target

The patients’ survival analysis is one of the major concerns to focus on certain gene to use as diagnosis tool or drug target. We have analyzed the relation between patients’ survival and *CDK4* expression (Figure 6). In both SARC (Figure 6a) and LGG (Figure 6b), patients’ survival is directly correlated with the higher expression of *CDK4*. This data corroborates with our previously shown expression profile of *CDK4* at different stage of SARC, LGG, and GBM (Figure 5). As *CDK4* expression is directly linked to the patients’ survival events, next we have compared the *CDK4* expression level in other 33 sub-types (Figure 3), where *CDK4* expression was detected, and found that *CDK4* expression is significantly higher in all of these subtypes compared to *TP53* (Figure 7). Altogether, the CDK4 expression profiling at 33 cancer sub-types and correlation with the patients’ survival has emphasized its importance as diagnostic tool and anticancer target, which we aimed based on our phytochemical screening (Figure 1, 2).

**Figure 6:**
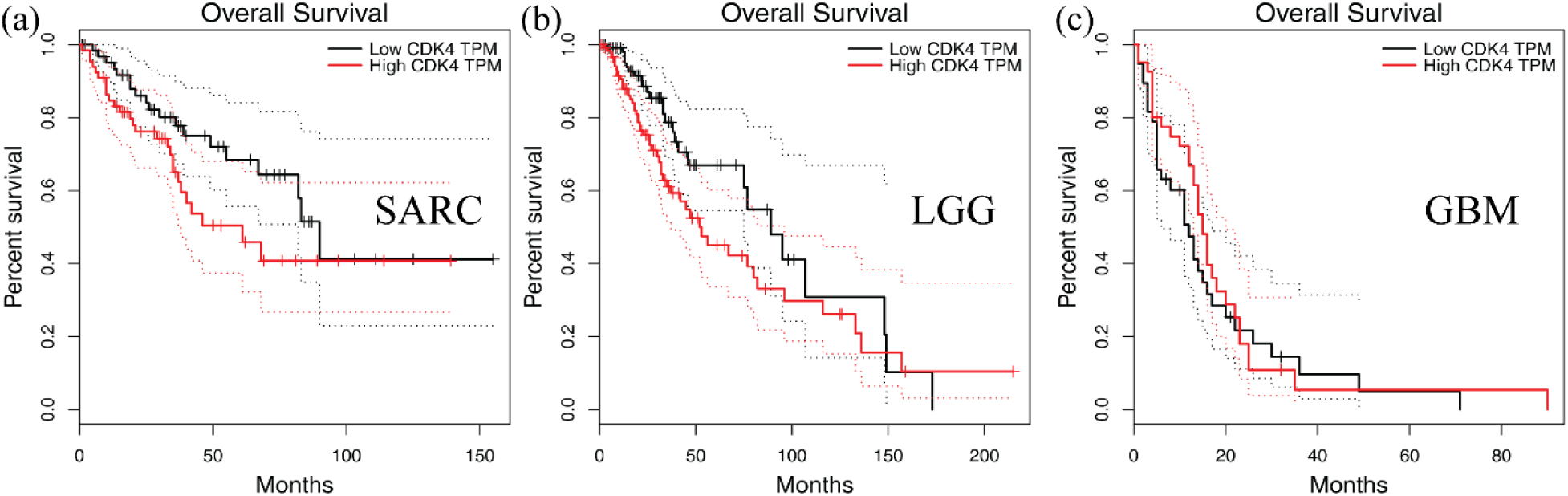
Relation between *CDK4* and survival rate of patients. *CDK4* expression level at different time points (months) of SARC (a), LGG (b), and GBM (c) patients.

**Figure 7:**
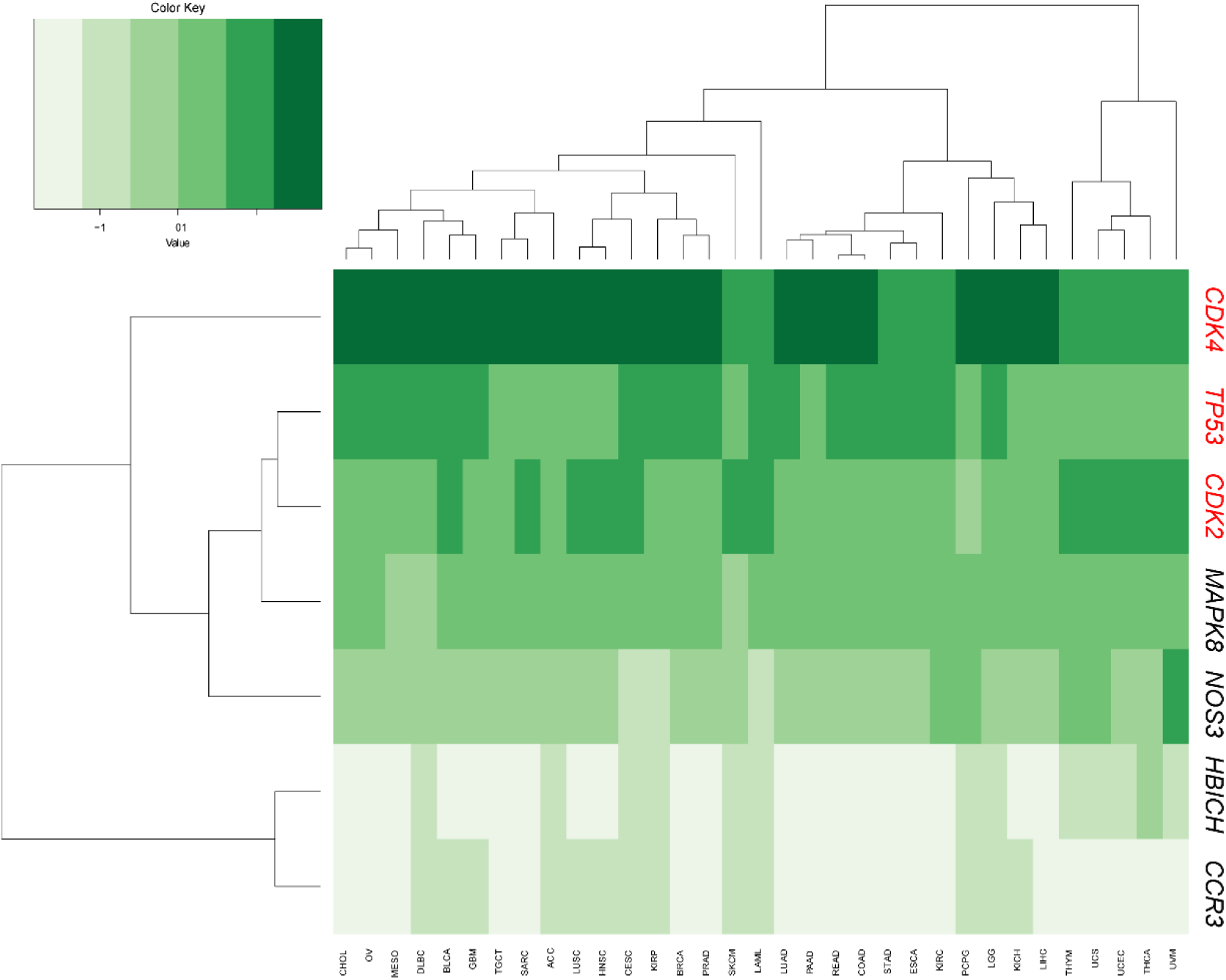
CDK4 express consistently in multiple cancers. Gene expression of CDK4, TP53, CDK2, MAPK8, HBICH, and CCR3 are compared in ACC (Adrenocortical carcinoma), BLCA (Bladder Urothelia Carcinoma), BRCA (Breast invasive carcinoma), CESC (Cervical squamous cell carcinoma and endocervical adenocarcinoma), CHOL (Cholangio carcinoma), COAD (Colon adenocarcinoma), DLBC (Lymphoid Neoplasm Diffuse Large B-cell Lymphoma), ESCA (Esophageal carcinoma), GBM (Glioblastoma multiforme), HNSC (Head and Neck squamous cell carcinoma), KICK (Kidney Chromophobe), KIRC (Kidney Renal clear cell carcinoma), KIRP (Kidney renal papillary cell carcinoma), LAML (Acute Myeloid Leukemia), LGG (Brain Lower Grade Glioma), LIHC (Liver hepatocellular carcinoma), LUAD (Lung adenocarcinoma), LUSC (Lung squamous cell carcinoma), MESO (Mesothelioma), OV (Ovarian serous cystadenocarcinoma), PAAD (Pancreatic adenocarcinoma), PCPG (Pheochromocytoma and Paraganglioma), PRAD (Prostate adenocarcinoma), READ (Rectum adenocarcinoma), SARC (Sarcoma), SKMC (Skin Cutaneous Melanoma), STAD (Stomach adenocarcinoma), TGCT (Testicular Germ Cell Tumors), THCA (Thyroid carcinoma), THYM (Thymoma), UCEC (Uterine Corpus Endometrial Carcinoma), UCS (Uterine Carcinosarcoma), UVM (Uveal Melanoma).

Additionally, we have identified the other genes (*METTL1, METTL21B, TSFM, OS9, MARCH9, TSPAN31*) which co-express along with *CDK4* (Figure 8). *CDK4* expression correlates to *METTL1* (Figure 8a), *METTL21B* (Figure 8b), *TSFM* (Figure 8c), *OS9* (Figure 8d), *MARCH9* (Figure 8e), *TSPAN31* (Figure 8f) with a R value 0.74, 0.61, 0.61, 0.57, 0.5, 0.41; respectively. This co-expression profiling is helpful to develop a set of genes along with *CDK4*, where mangiferin effect will be demonstrated by their expression level.

**Figure 8:**
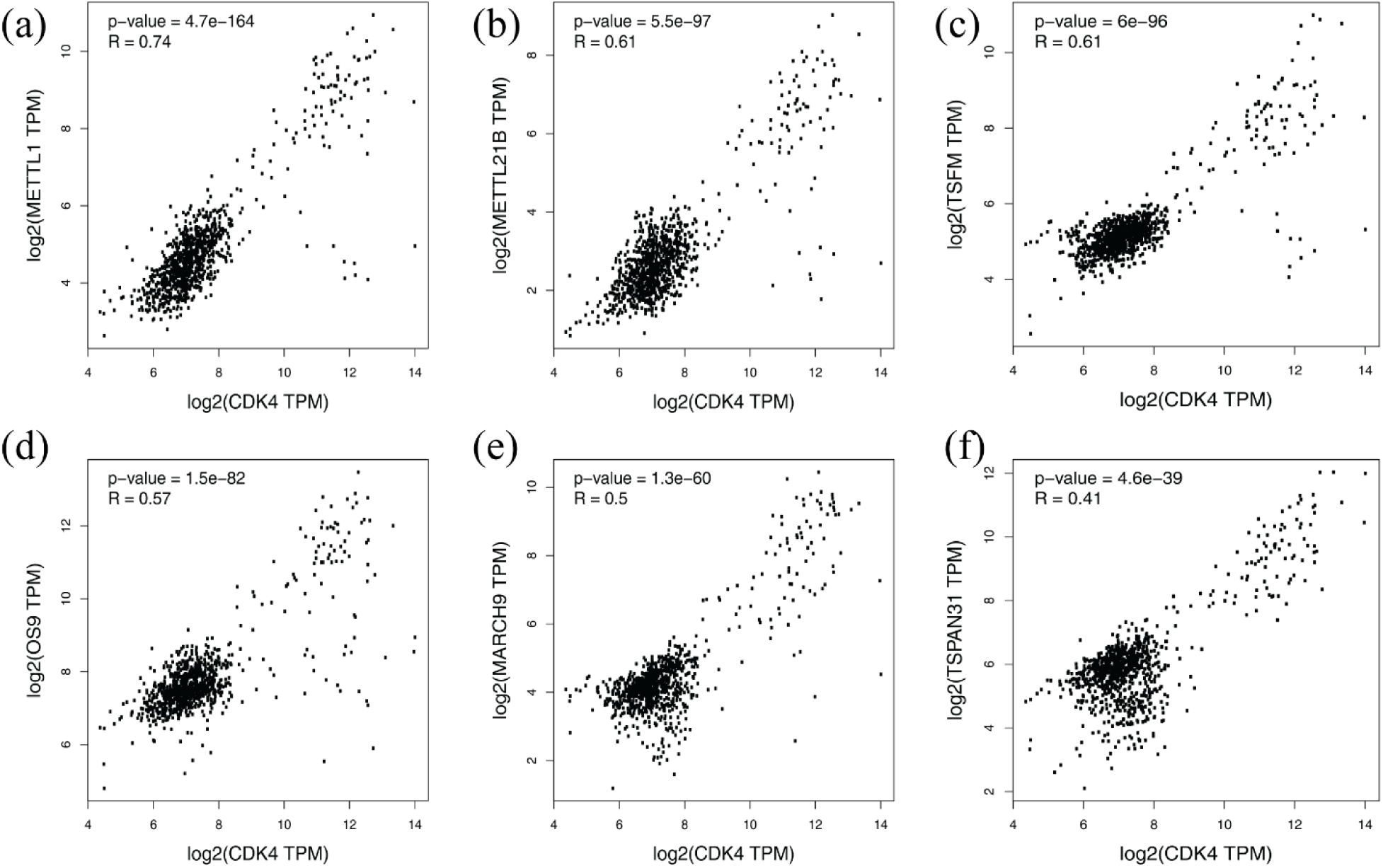
Co-expression analysis between *CDK4* and closely related genes. Co-expression of *CDK4* with *METTL1* (a), *METTL21B* (b), *TSFM* (c), *OS9* (d), *MARCH9* (e), and *TSPAN31* (f). Log2 value of TPM (Transcripts per million) from each gene was considered correlation analysis, P-and R value are embedded in each graph.

## Discussion

In the recent years, due to the effort of Moonshot^SM^ project and an increase amount of interest among local scientists to develop medicinal plant databases (Ashraf et al. 2014; Brito and Brito 1993; Babu et al. 2006; Mohammed et al. 2010; Mohanraj, Karthikeyan, Vivek-Ananth, et al. 2018), the trend of using phytochemical as drug targets in reviving. Our endeavor started with the MPDB1.0 (Ashraf et al. 2014) and extended through MPDB2.0. This database provides a huge resource to start the screening. From our screening, we have identified mangiferin. This simple compound is not only available in our food items, but also used in traditional medicinal practice. It has been known as treatment for diabetes, infection, and cancer (Kavitha et al. 2013; Tolosa et al. 2013). Natural medicine, Vimang®, is produced from *Mangifera indica* extracts and used for anti-inflammatory phytomedicine (Rajendran et al., 2014). Previous studies suggest that mangiferin regulates Mitogen Activated Protein Kinase pathway and progression of G2/M phase of cell cycle (Lv et al. 2013). Consistent with that, we have identified MAPK8, CDK2, and CDK4 in our list of interacting proteins (Figure 2). Fundamentally, this study identified one of the cell cycle regulators, CDK4, which was predicted in other complementary studies.

Interestingly, CDKs are known to be upregulated during cancer and CDK inhibitors are used as anticancer drug trial (Goel et al. 2018; J. H. Jeong et al. 2009; Yin et al. 2018; Malumbres and Barbacid 2009; Vijayaraghavan et al. 2018). As a result, finding mangiferin targeting CDK4 as anticancer drug target has validated our approach (Figure 2). Even in five years back, it was not straight forward to test our hypothesis and screen active compounds from MPDB1.0 for their anticancer activity. During the last few years, availability of cancer related RNA sequencing datasets such as TCGA and GTEx has made it possible (Consortium 2015; Lonsdale et al. 2013; Weinstein et al. 2013). Our analysis to link the mangiferin with CDK4 was accelerated by using this large-scale RNA sequencing data. Our analysis of *CDK4* expression in the different cancer sub-types has opened the new door to develop *CDK4* as a cancer diagnostic tool as it is expressed prominently in multiple cancer sub-types throughout different stages and correlate with patients’ survival (Figure 3, 4, 5, 6, 7). It has also emphasized on cyclin-dependent kinase as a general target to study against mangiferin and its derivative in near future.

Based on the current study, we have developed the hypothesis that mangiferin is targeting cell cycle regulator CDK4, which is upregulated in majority of cancer events, and mango tree (*Mangifera indica*) derived extract mangiferin can be used as inhibitor of CDK4 to suppress its expression and used as potential anticancer drug (Figure 9).

**Figure 9:**
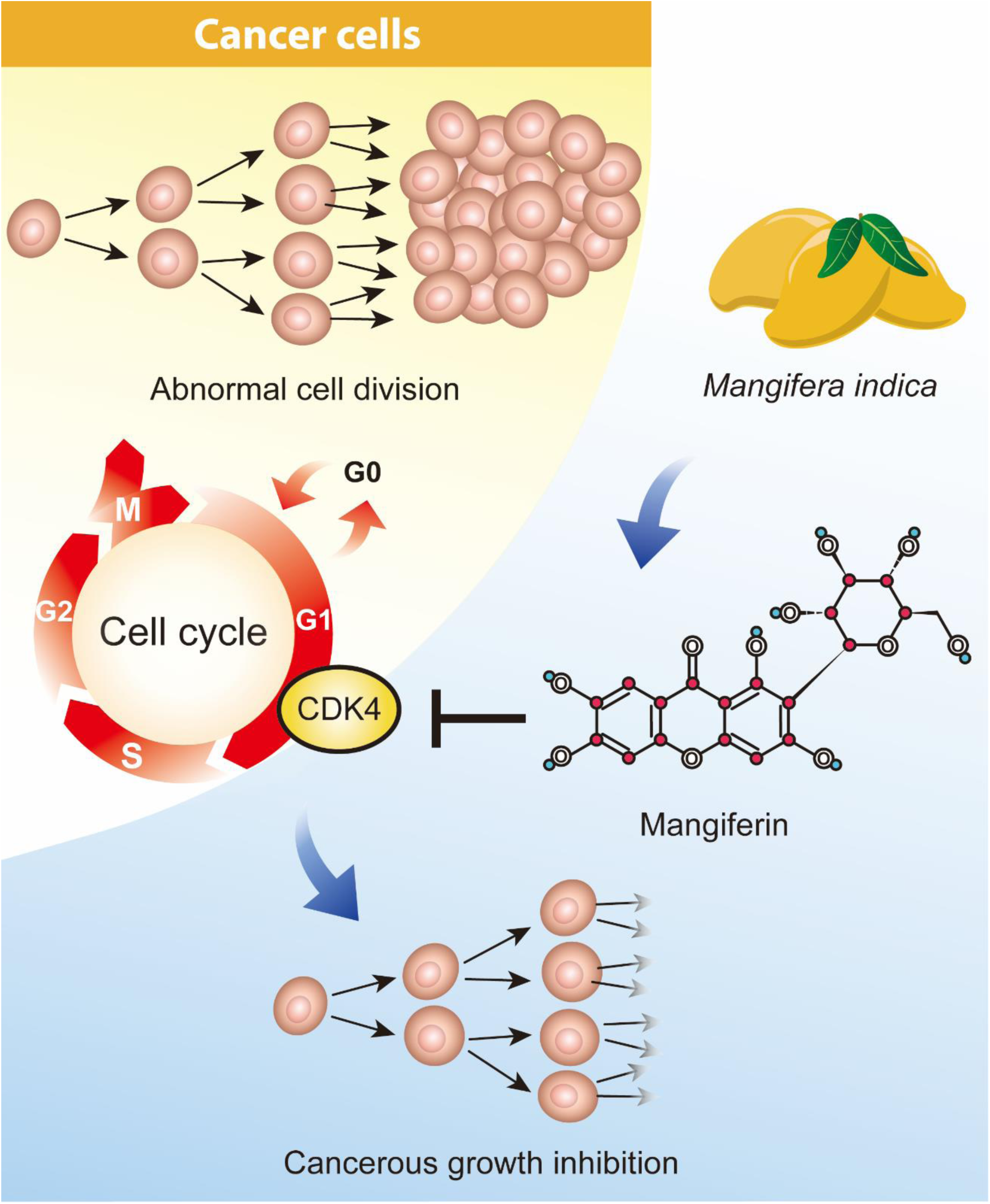
Proposed working model based on the current study.

## Conclusions

Mangiferin, a mango tree (*Mangifera indica*) derived extract, which is inexpensive and easily available is one of the potential CDK4 inhibitors. As the extraction procedure of purified mangiferin and commercially purified mangiferin are available, it will help us to quickly move forward to test the hypothesis whether mangiferin can inhibit CDK4 to regulate cancerous growth. Fundamentally, this study ushered the way of developing affordable anticancer drugs.

## Conflicts of Interest

The authors declare no conflicts of interest.

## Authors’ contributions

R.K.C., N.H., and M.S.R. collected the medicinal plant and their active compounds’ information. R.K.C. and N.H. studied the interacting proteins for active compounds. S.A. executed the molecular docking. M.A.A. did the large-scale gene expression analysis. S.S. did the gene expression analysis for the heatmap. T.A.F. is responsible for the curation of database. A.A.T. and F.M. contributed from the MPDB1.0 team. M.K.H. supervised R.K.C. and N.H. and M.A.S supervised M.S.R. for this project. Manuscript draft was written by M.K.H. and S.S. E.H.A. provided his expert opinion to improve the manuscript. M.K.H., M.A.S., and M.A.A. coordinated the project. M.A.A. has adopted the project idea, designed experiments, prepared final figures, and wrote the manuscript.

## Data availability

Data used in this study or produced is either available from the public domain and mentioned here in the manuscript. Apart from that, authors always welcome to share the data required for reviewers and other researchers.

## Funding statement

This project is currently not funded.

## Acknowledgement

Authors would like to thank researchers around the globe for their relentless effort to put together the information of medicinal plants.

## References

Al-kalaldeh, Jelnar Z, Rana Abu-dahab, and Fatma U Afifi. 2010. “Volatile Oil Composition and Antiproliferative Activity of Laurus Nobilis, Origanum Syriacum, Origanum Vulgare, and Salvia Triloba against Human Breast Adenocarcinoma Cells.” Nutrition Research 30 (4): 271–78. https://doi.org/10.1016/j.nutres.2010.04.001.

Altunkaya, Ali, Chunxiao Bi, Anthony R Bradley, Peter W Rose, Andreas Prli, H Christie, Luigi Di Costanzo, et al. 2017. “The RCSB Protein Data Bank: Integrative View of Protein, Gene and 3D Structural Information” 45 (October 2016): 271–81. https://doi.org/10.1093/nar/gkw1000.

Ashraf, Mohammad Arif, Achia Khatun, Tanzila Sharmin, Faraid Mobin, Arifur Rahman Tanu, Toufique Morshed, Tawkir Ahmad Fakir, Rifat Ara Begum, and A H M Nurun Nabi. 2014. “MPDB 1.0: A Medicinal Plant Database of Bangladesh.” Bioinformation 10 (6): 384.

Babu, Padavala Ajay, Gadde Suneetha, Radha Boddepalli, Vedurupaka Vasantha Lakshmi, Talluru Sudha Rani, Yellapu RamBabu, and Kolli Srinivas. 2006. “A Database of 389 Medicinal Plants for Diabetes.” Bioinformation 1 (4): 130.

Balunas, Marcy J, and A Douglas Kinghorn. 2005. “Drug Discovery from Medicinal Plants” 78: 431–41. https://doi.org/10.1016/j.lfs.2005.09.012.

Bray, Freddie, Jacques Ferlay, and Isabelle Soerjomataram. 2018. “Global Cancer Statistics 2018: GLOBOCAN Estimates of Incidence and Mortality Worldwide for 36 Cancers in 185 Countries,” 394–424. https://doi.org/10.3322/caac.21492.

Bridgeman, Christopher J., Thuy Uyen Nguyen, and Vipuil Kishore. 2018. “Anticancer Efficacy of Tannic Acid Is Dependent on the Stiffness of the Underlying Matrix.” Journal of Biomaterials Science, Polymer Edition 29 (4): 412–27. https://doi.org/10.1080/09205063.2017.1421349.

Brito, Alba R M Souza, and Antonio A Souza Brito. 1993. “Forty Years of Brazilian Medicinal Plant Research.” Journal of Ethnopharmacology 39 (1): 53–67.

Butler, Mark S. 2004. “The Role of Natural Product Chemistry in Drug Discovery †,” 2141–53.

Consortium, GTEx. 2015. “The Genotype-Tissue Expression (GTEx) Pilot Analysis: Multitissue Gene Regulation in Humans.” Science 348 (6235): 648–60.

Cook, Matthew T. 2018. “Mechanism of Metastasis Suppression by Luteolin in Breast Cancer.” Breast Cancer: Targets and Therapy 10: 89.

Cragg, Gordon M, and David J Newman. 2005. “Plants as a Source of Anti-Cancer Agents” 100: 72–79. https://doi.org/10.1016/j.jep.2005.05.011.

Dallakyan, Sargis, and Arthur J Olson. 2015. “Chapter 19 Small-Molecule Library Screening by Docking with PyRx” 1263: 243–50. https://doi.org/10.1007/978-1-4939-2269-7.

Desai, Avni G, Ghulam N Qazi, Ramesh K Ganju, Mahmoud El-tamer, Jaswant Singh, Ajit K Saxena, Yashbir S Bedi, Subhash C Taneja, and Hari K Bhat. 2008. “Medicinal Plants and Cancer Chemoprevention,” 581–91.

Gaikwad, Jitendra, Varun Khanna, Subramanyam Vemulpad, Joanne Jamie, Jim Kohen, and Shoba Ranganathan. 2008. “CMKb: A Web-Based Prototype for Integrating Australian Aboriginal Customary Medicinal Plant Knowledge” 8: 1–8. https://doi.org/10.1186/1471-2105-9-S12-S25.

Goel, Shom, Molly J DeCristo, Sandra S McAllister, and Jean J Zhao. 2018. “CDK4/6 Inhibition in Cancer: Beyond Cell Cycle Arrest.” Trends in Cell Biology 28 (11): 911–25.

Jeong, Jae-Hoon, Jee Young An, Yong Tae Kwon, Juong G Rhee, and Yong J Lee. 2009. “Effects of Low Dose Quercetin: Cancer Cell-specific Inhibition of Cell Cycle Progression.” Journal of Cellular Biochemistry 106 (1): 73–82.

Jeong, Jae Hoon, Jee Young An, Yong Tae Kwon, Juong G. Rhee, and Yong J. Lee. 2009. “Effects of Low Dose Quercetin: Cancer Cell-Specific Inhibition of Cell Cycle Progression.” Journal of Cellular Biochemistry 106 (1): 73–82. https://doi.org/10.1002/jcb.21977.

Kavitha, Mani, Jagatheesan Nataraj, Musthafa Mohammed Essa, Mushtaq A Memon, and Thamilarasan Manivasagam. 2013. “Mangiferin Attenuates MPTP Induced Dopaminergic Neurodegeneration and Improves Motor Impairment, Redox Balance and Bcl-2/Bax Expression in Experimental Parkinson’s Disease Mice.” Chemico-Biological Interactions 206 (2): 239–47.

Khan, F, P Pandey, T K Upadhyay, A Jafri, N K Jha, R Mishra, and V Singh. 2019. “Anti-Cancerous Effect of Rutin Against HPV-C33A Cervical Cancer Cells via G0/G1 Cell Cycle Arrest and Apoptotic Induction.” Endocrine, Metabolic & Immune Disorders Drug Targets.

Khatun, Amina, Mahmudur Rahman, Tania Haque, Mahfizur Rahman, Mahfuja Akter, Subarna Akter, and Afrin Jhumur. 2014. “Cytotoxicity Potentials of Eleven Bangladeshi Medicinal Plants” 2014.

Kim, Sunghwan, Paul A Thiessen, Evan E Bolton, Jie Chen, Gang Fu, Asta Gindulyte, Lianyi Han, et al. 2016. “PubChem Substance and Compound Databases” 44 (September 2015): 1202–13. https://doi.org/10.1093/nar/gkv951.

Kim, Yonghyo, Hye Jin Jung, and Ho Jeong Kwon. 2012. “A Natural Small Molecule Voacangine Inhibits Angiogenesis Both in Vitro and in Vivo.” Biochemical and Biophysical Research Communications 417 (1): 330–34.

Kuhn, Michael, Christian Von Mering, Monica Campillos, and Lars Juhl Jensen. 2008. “STITCH: Interaction Networks of Chemicals and Proteins” 36 (December 2007): 684–88. https://doi.org/10.1093/nar/gkm795.

Liu, Zuojia, Dan Li, Lijun Yu, and Fenglan Niu. 2012. “Gallic Acid as a Cancer-Selective Agent Induces Apoptosis in Pancreatic Cancer Cells.” Chemotherapy 58 (3): 185–94.

Lonsdale, John, Jeffrey Thomas, Mike Salvatore, Rebecca Phillips, Edmund Lo, Saboor Shad, Richard Hasz, Gary Walters, Fernando Garcia, and Nancy Young. 2013. “The Genotype-Tissue Expression (GTEx) Project.” Nature Genetics 45 (6): 580.

Lv, Jianzhen, Zijie Wang, Li Zhang, Hai-Lian Wang, Yuande Liu, Chunyang Li, Jiagang Deng, Yi-Wang, and Jin-Ku Bao. 2013. “Mangiferin Induces Apoptosis and Cell Cycle Arrest in MCF-7 Cells Both in Vitro and in Vivo.” JOURNAL OF ANIMAL AND VETERINARY ADVANCES 12 (3): 352–59.

Malumbres, Marcos, and Mariano Barbacid. 2009. “Cell Cycle, CDKs and Cancer: A Changing Paradigm” 9 (mArCH). https://doi.org/10.1038/nrc2602.

Mohammed, Rahmatullah, Ferdausi Dilara, Mollik Md Ariful Haque, Jahan Rownak, Chowdhury Majedul H., and Haque Mozammel Wahid. 2010. “A SURVEY OF MEDICINAL PLANTS USED BY KAVIRAJES OF CHALNA AREA, KHULNA” 7: 91–97.

Mohanraj, Karthikeyan, Bagavathy Shanmugam Karthikeyan, R P Bharath Chand, S R Aparna, Pattulingam Mangalapandi, and Areejit Samal. 2018. “OPEN IMPPAT: A Curated Database of Indian Medicinal Plants, Phytochemistry And Therapeutics.” Scientific Reports, no. February: 1–17. https://doi.org/10.1038/s41598-018-22631-z.

Mohanraj, Karthikeyan, Bagavathy Shanmugam Karthikeyan, R P Vivek-Ananth, R P Bharath Chand, S R Aparna, Pattulingam Mangalapandi, and Areejit Samal. 2018. “IMPPAT: A Curated Database of I Ndian M Edicinal P Lants, P Hytochemistry A Nd T Herapeutics.” Scientific Reports 8 (1): 4329.

Mukherjee, Ashis K, Sourav Basu, Nabanita Sarkar, and Anil C Ghosh. 2001. “Advances in Cancer Therapy with Plant Based Natural Products,” 1467–86.

Norman, R. Farnsworth, Akerele Olaywola, Bingel Audrey S., D. Soejarto Djaja, and Guo Zhengang. 1985. “Medicinal Plants in Therapy *.”

Noronha, Vanita, Tata Memorial Centre, Arif Jamshed, Shaukat Khanum, Memorial Cancer, Kumar Prabhash, and Tata Memorial Centre. 2012. “A Fresh Look at Oncology Facts on South Central Asia and SAARC Countries,” no. May 2014. https://doi.org/10.4103/2278-330X.96489.

Núñez Selles, Alberto J, Maria Daglia, and Luca Rastrelli. 2016. “The Potential Role of Mangiferin in Cancer Treatment through Its Immunomodulatory, Anti-angiogenic, Apoptopic, and Gene Regulatory Effects.” Biofactors 42 (5): 475–91.

Oleg, Trott, and Olson Arthur J. 2011. “NIH Public Access” 31 (2): 455–61. https://doi.org/10.1002/jcc.21334.AutoDock.

Peng, Yulin, Yan Wang, Ning Tang, Dongdong Sun, Yulong Lan, Zhenlong Yu, Xinyu Zhao, Lei Feng, Baojing Zhang, and Lingling Jin. 2018. “Andrographolide Inhibits Breast Cancer through Suppressing COX-2 Expression and Angiogenesis via Inactivation of P300 Signaling and VEGF Pathway.” Journal of Experimental & Clinical Cancer Research 37 (1): 248.

Rajendran, Peramaiyan, Thangavel Jayakumar, Ikuo Nishigaki, Yutaka Nishigaki, Jayabal Vetriselvi, and Dhanapal Sakthisekaran. n.d. “Immunomodulatory Effect of Mangiferin in Experimental Animals with Benzo (a) Pyrene-Induced Lung Carcinogenesis” 9 (2): 61–67.

Saddala, Madhu Sudhana, Pradeep Kiran, Jangampalli Adi, and Usha Rani A. 2016. “In Silico Drug Design and Molecular Docking Studies of Potent Inhibitors against Cathepsin-L (Ctsl) for Sars Disease” 4 (2): 2–5.

Singh, Anjana, Shyam Singh Chauhan, and Vishwas Tripathi. 2018. “Quinic Acid Attenuates Oral Cancer Cell Proliferation by Downregulating Cyclin D1 Expression and Akt Signaling.” Pharmacognosy Magazine 14 (55): 14.

Sousa, Filipe, Pedro Alexandrino Fernandes, and Maria Joa. 2006. “Protein – Ligand Docking: Current Status and Future” 26 (February): 15–26. https://doi.org/10.1002/prot.

Tang, Zefang, Boxi Kang, Chenwei Li, Tianxiang Chen, and Zemin Zhang. 2019. “GEPIA2: An Enhanced Web Server for Large-Scale Expression Profiling and Interactive Analysis.” Nucleic Acids Research.

Tang, Zefang, Chenwei Li, Boxi Kang, Ge Gao, Cheng Li, and Zemin Zhang. 2017. “GEPIA: A Web Server for Cancer and Normal Gene Expression Profiling and Interactive Analyses.” Nucleic Acids Research 45 (W1): W98–102.

Tolosa, Laia, Idania Rodeiro, M Teresa Donato, José A Herrera, René Delgado, José V Castell, and M José Gómez-Lechón. 2013. “Multiparametric Evaluation of the Cytoprotective Effect of the M Angifera Indica L. Stem Bark Extract and Mangiferin in HepG2 Cells.” Journal of Pharmacy and Pharmacology 65 (7): 1073–82.

Uddin, A F M Kamal, Zohora Jameela Khan, Johirul Islam, and Mahmud Am. 2016. “Cancer Care Scenario in Bangladesh” 2 (2): 102–4. https://doi.org/10.4103/2278-330X.110510.

Vijayaraghavan, Smruthi, Stacy Moulder, Khandan Keyomarsi, and Rachel M Layman. 2018. “Inhibiting CDK in Cancer Therapy: Current Evidence and Future Directions.” Targeted Oncology 13 (1): 21–38.

Wang, Yanli, Evan Bolton, Svetlana Dracheva, Karen Karapetyan, Benjamin A Shoemaker, Tugba O Suzek, Jiyao Wang, Jewen Xiao, Jian Zhang, and Stephen H Bryant. 2010. “An Overview of the PubChem BioAssay Resource” 38 (November 2009): 255–66. https://doi.org/10.1093/nar/gkp965.

Wang, Yanli, Jewen Xiao, Tugba O Suzek, Jian Zhang, Jiyao Wang, and Stephen H Bryant. 2009. “PubChem: A Public Information System for Analyzing Bioactivities of Small Molecules” 37 (June): 623–33. https://doi.org/10.1093/nar/gkp456.

Wang, Yanli, Jewen Xiao, Tugba O Suzek, Jian Zhang, Jiyao Wang, Zhigang Zhou, Lianyi Han, et al. 2012. “PubChem‘s BioAssay Database” 40 (December 2011). https://doi.org/10.1093/nar/gkr1132.

Weinstein, John N, Eric A Collisson, Gordon B Mills, Kenna R Mills Shaw, Brad A Ozenberger, Kyle Ellrott, Ilya Shmulevich, Chris Sander, Joshua M Stuart, and Cancer Genome Atlas Research Network. 2013. “The Cancer Genome Atlas Pan-Cancer Analysis Project.” Nature Genetics 45 (10): 1113.

Yan, Xiaohui, Miao Qi, Pengfei Li, Yihong Zhan, and Huanjie Shao. 2017. “Apigenin in Cancer Therapy: Anti - Cancer Effects and Mechanisms of Action.” Cell & Bioscience, 1–16. https://doi.org/10.1186/s13578-017-0179-x.

Yin, Xifeng, Jun Yu, Yang Zhou, Chengyue Wang, Zhimin Jiao, and Zhounan Qian. 2018. “Identification of CDK2 as a Novel Target in Treatment of Prostate Cancer.”

